# Chlamydial YAP activation in host endocervical epithelial cells mediates pro-fibrotic paracrine stimulation of fibroblasts

**DOI:** 10.1101/2023.05.30.542940

**Authors:** Liam Caven, Rey Carabeo

## Abstract

Infection of the female genital tract by *Chlamydia trachomatis* can produce severe fibrotic sequelae, including tubal factor infertility and ectopic pregnancy. While infection demonstrably mediates a pro-fibrotic response in host cells, it remains unclear if intrinsic properties of the upper genital tract exacerbate chlamydial fibrosis. The relatively sterile environment of the upper genital tract is primed for a pro-inflammatory response to infection, potentially enhancing fibrosis - however, subclinical *C. trachomatis* infections still develop fibrosis-related sequelae. Here, we compare infection-associated and steady-state gene expression of primary human cervical and vaginal epithelial cells. In the former, we observe enhanced baseline expression and infection-mediated induction of fibrosis-associated signal factors (e.g. *TGFA*, *IL6*, *IL8*, *IL20*), implying predisposition to *Chlamydia*-associated pro-fibrotic signaling. Transcription factor enrichment analysis identified regulatory targets of YAP, a transcriptional cofactor induced by infection of cervical epithelial cells, but not vaginal epithelial cells. YAP target genes induced by infection include secreted fibroblast-activating signal factors; therefore, we developed an *in vitro* model involving coculture of infected endocervical epithelial cells with uninfected fibroblasts. Coculture enhanced fibroblast expression of type I collagen, as well as prompting reproducible (albeit statistically insignificant) induction of α-smooth muscle actin. Fibroblast collagen induction was sensitive to siRNA-mediated YAP knockdown in infected epithelial cells, implicating chlamydial YAP activation in this effect. Collectively, our results present a novel mechanism of fibrosis initiated by *Chlamydia,* wherein infection-mediated induction of host YAP facilitates pro-fibrotic intercellular communication. Chlamydial YAP activation in cervical epithelial cells is thus a determinant of this tissue’s susceptibility to fibrosis.

**Importance:** Chronic or repeated infection of the female upper genital tract by *C. trachomatis* can lead to severe fibrotic sequelae, including tubal factor infertility and ectopic pregnancy. However, the molecular mechanisms underlying this effect are unclear. In this report, we define a transcriptional program specific to *C. trachomatis* infection of the upper genital tract, identifying tissue-specific induction of host YAP – a pro-fibrotic transcriptional cofactor – as a potential driver of infection-mediated fibrotic gene expression. Further, we show that infected endocervical epithelial cells stimulate collagen production by fibroblasts, and implicate chlamydial induction of YAP in this effect. Our results define a mechanism by which infection mediates tissue-level fibrotic pathology via paracrine signaling, and identify YAP as a potential therapeutic target for prevention of *Chlamydia*-associated scarring of the female genital tract.

## Introduction

The Gram-negative, obligate intracellular pathogen C*hlamydia trachomatis* is responsible for the most common bacterial sexually transmitted infection worldwide. Critically, it is estimated that up to 80% of *C. trachomatis* infections of the female genital tract are asymptomatic ^1^. The relatively poor state of *C. trachomatis* surveillance is of particular concern to public health, given the severe complications associated with chronic or repeated infection. In females, dissemination of the infection into the upper genital tract can lead to pelvic inflammatory disease (PID) ^2, 3^. Additionally, chlamydial infection of the Fallopian tubes is associated with progressive scarring that can block descension of the ovum, thereby resulting in ectopic pregnancy or tubal factor infertility (TIF) ^4, 5^. A retrospective study of reproductive complications following screening for *C. trachomatis* in Danish women found that repeat infections increased the risk of PID by 20% ^6^. Similarly, a study modeling the risk of specific infection-associated sequelae using epidemiological data from the United Kingdom estimated that up to 29% of all TIF cases are attributable to *C. trachomatis* infection ^7^. Given these data, it is unsurprising that the study of chlamydial pathogenesis and the mechanisms underlying *Chlamydia*-associated fibrotic sequelae comprise topics of ongoing investigation.

Recent work suggests that *C. trachomatis* induces pro-fibrotic signaling by host epithelial cells. *Chlamydia* infection has been shown to induce signal factors associated with myofibroblast activation and wound healing, such as IL-6, IL-11, EGF, and CTGF ^8–12^. The related species *C. pneumoniae* exhibits similar induction of pro-fibrotic signaling ‒ including production of TGF-β1, a consistent stimulus of myofibroblast differentiation ^13–15^. Additionally, infection-mediated induction of the epithelial-to-mesenchymal transition (EMT) in host epithelial cells has been consistently reported over the past decade, including upregulation of mesenchymal markers such as N-cadherin, MMP9, and fibronectin, as well as the pro-EMT regulators ZEB1/2, SNAIL/SLUG, and thrombospondin 1 ^16–18^. Epithelial cells undergoing EMT transdifferentiate into a myofibroblast-like phenotype, subsequently remodeling the extracellular matrix (ECM) to drive scar formation ^19^. Accordingly, it has been observed that *Chlamydia* infection induces host expression of genes associated with ECM remodeling, including ECM components (collagens, laminins), and ECM-modifying matrix metalloproteases ^20, 21^. Taken together, these data lend credence to a model of chlamydial fibrosis enhanced by, but not necessarily dependent upon immune cell recruitment to the site of infection.

Chlamydial induction of EMT has been associated with deposition of fibrillar collagen – a common hallmark of scar formation – however, the mechanisms underlying *Chlamydia*-induced fibrosis remain incompletely characterized. In particular, it is unclear to what extent the pro-fibrotic signaling of infected epithelial cells influences other cell types not directly accessible for infection by *Chlamydia*, such as tissue-resident fibroblasts. Importantly, signal factors produced by *Chlamydia*-infected epithelial cells have been shown in other contexts to promote fibroblast differentiation into contractile and collagen-producing myofibroblasts ^22–25^. Given that aberrant fibroblast activation has been linked to idiopathic pulmonary fibrosis and other forms of scarring-associated disease ^26^, an assessment of the effect of *Chlamydia*-associated signaling on this cell type may provide key insight into the mechanisms underlying chlamydial fibrosis.

While the terminally differentiated epithelial cells of the *stratum corneum* are considered a poor host for *C. trachomatis*, given their lack of mitochondria and attendant metabolic inactivity ^27, 28^, a recent report from our laboratory indicated that undifferentiated epithelial cells from this tissue can support *C. trachomatis* serovars D and L2 in a three-dimensional organotypic model of infection ^29^. Accordingly, a study comparing *C. trachomatis* serovar E infection of squamous ectocervical epithelial cells and endometrial epithelial cells cultured *in vitro* reported viable infection of the former, albeit at reduced efficiency ^30^. In combination with more recent reports of early passage immortalized human vaginal epithelial cells supporting infection of genital (serovar D) and LGV (serovar L2) *C. trachomatis* biovars ^27^, these data collectively illustrate the pathogen’s capacity to infect transitional or undifferentiated vaginal epithelial cells. Critically, fibrotic pathology is rarely reported in *Chlamydia*-infected vaginal epithelium – suggesting that intrinsic differences between cervical and vaginal epithelial cells may contribute to *Chlamydia*-associated scarring of the upper genital tract.

To determine if infection of a specific cell type is associated with pro-fibrotic gene expression, we have examined the host response to *C. trachomatis* in primary human cervical epithelial cells (HCECs) and primary human vaginal epithelial cells (HVEs). We observe both steady-state and infection-associated induction of pro-inflammatory and pro-fibrotic signaling in the former, including increased expression of *TGFA*, *IL6*, *IL8*, and *IL20*. Transcription factor enrichment analysis of this gene set implicated a broad portfolio of known regulatory targets of YAP, a pro-fibrotic transcriptional cofactor. Infection promoted YAP activation in cervical epithelial cells, but not vaginal epithelial cells – suggesting YAP may contribute to pro-fibrotic gene expression specific to the upper genital tract. To assess the downstream effects of pro-fibrotic/pro-inflammatory signaling by infected cells, we have developed an *in vitro* model of infection wherein *Chlamydia*-infected, immortalized endocervical epithelial cells (End1s) are cocultured with uninfected uterine fibroblasts (KCO2s). We show that that fibroblasts cocultured with infected epithelial cells exhibit significantly increased expression of collagen I – a phenotype sensitive to siRNA-mediated knockdown of YAP in host epithelial cells. Taken together, our results define a novel means by which infection promotes fibrotic gene and protein expression, dependent upon host YAP activity and intercellular communication.

## Results

### Cervical epithelial cells exhibit intrinsic pro-inflammatory and pro-EMT gene expression relative to vaginal epithelial cells

To identify gene expression potentially driving fibrotic pathologies in the epithelial tissues of the upper genital tract, we compared the transcriptomes of mock and *Chlamydia*-infected cervical epithelial cells to that of infected vaginal epithelial cells. Primary human cervical epithelial cells (HCECs) and vaginal epithelial cells (HVEs) were either mock-infected or infected with the anogenital *C. trachomatis* serovar L2, then harvested for polyadenylated RNA for bulk RNA-sequencing at 24 hours post-infection (hpi). Comparison of the *Chlamydia*-infected HCEC and HVE transcriptomes identified 7354 genes differentially expressed (FDRP ≤ 0.05) in infected cervical cells relative to infected vaginal cells (Figure 1A, Supplemental Table 1). Further characterization of this gene set was performed using the ranked functional enrichment analysis functions of the STRING database ^31, 32^. *Chlamydia*-infected HCECs exhibited increased expression of genes associated with pro-inflammatory signaling, with GO-BP (gene ontology, biological process) terms associated with a type-I interferon response, leukocyte chemotaxis, and complement activation showing significant enrichment (Table 1). Intriguingly, the infected HCEC transcriptome also exhibited enrichment of GO-BP terms associated with keratinocyte and epidermal differentiation (Table 1). Recent reports have also shown *Chlamydia* can induce an epithelial-to-mesenchymal transition (EMT) in endocervical epithelial cells – a pro-fibrotic transdifferentiation event associated with loss of cytokeratin expression ^16, 33, 34^. Accordingly, we observe significantly increased expression of EMT-regulating transcription factors such as *SNAIL/SNAI1* (FC: 7.35) and *TWIST1* (FC: 11.37) in infected HCECs relative to the infected HVE transcriptome. These data collectively suggest that cervical epithelial cells may be more susceptible than vaginal epithelial cells to infection-associated EMT.

**Figure 1.**
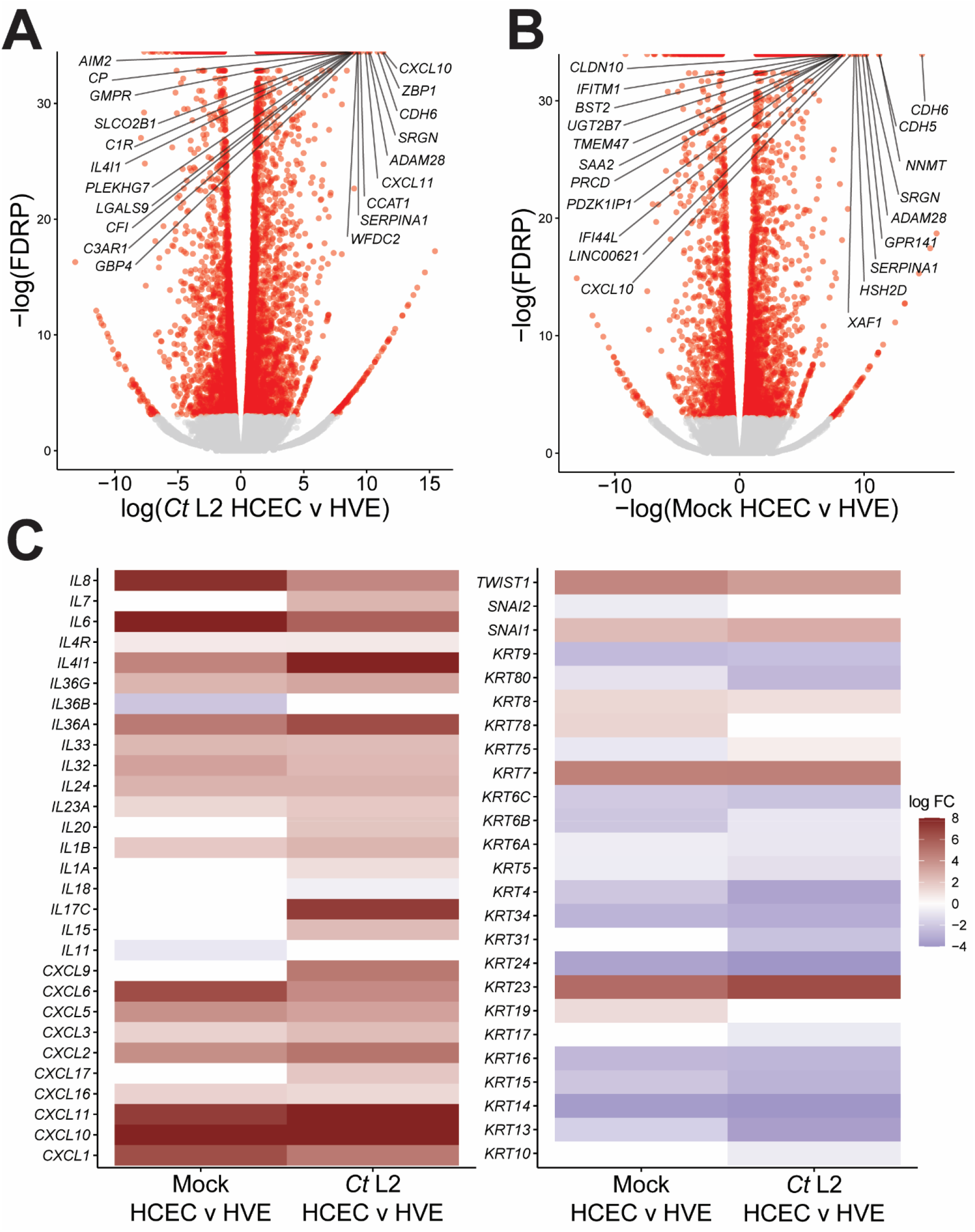
Cervical epithelial cells exhibit intrinsic pro-inflammatory and pro-EMT gene expression relative to vaginal epithelial cells. **(A, B)** Volcano plot of differential gene expression of *Chlamydia trachomatis* serovar L2-infected **(A)** or mock-infected **(B)** primary human cervical epithelial cells (HCECs) relative to equivalent human vaginal epithelial cells (HVEs) at 24 hpi (see also Supplementary Table 1). n = 3, with a minimum of 3×10^7^ unstranded single reads per replicate with a mean length of 150 bp. All fold changes are relative to the mock-infected control; red dots: false discovery rate p-value (FDRP) ≤ 0.05, labels: top 20 genes with lowest FDRP values. **(C)** Heatmap of cytokine and EMT-associated gene expression in mock-infected (left columns) and *C. trachomatis* serovar L2-infected (right columns) HCECs. All fold changes are relative to an equivalent infection in HVEs; only genes differentially expressed (FDRP ≤ 0.05) in at least one comparison are shown.

**Table 1:** GO Biological Process terms significantly enriched (FDR ≤ 0.05) in comparison of *C. trachomatis* serovar L2-infected primary human endocervical epithelial cells (HCECs) to primary human vaginal epithelial cells (HVEs) at 24 hours post-infection via bulk RNA-sequencing. Induction/repression of each term is relative to the HVE infection.

| **<u>GO-BP Term Description</u>** | **<u>Genes Mapped</u>** | **<u>Enrichment Score</u>** | **<u>Modulation</u>** | **<u>False Discovery Rate</u>** |
| --- | --- | --- | --- | --- |
| Ribonucleoprotein complex biogenesis | 166 | 0.81 | Repressed | 3.66E-23 |
| Ribosome biogenesis | 121 | 0.89 | Repressed | 2.02E-20 |
| ncRNA metabolic process | 171 | 0.70 | Repressed | 1.74E-19 |
| ncRNA processing | 136 | 0.80 | Repressed | 2.28E-18 |
| rRNA processing | 95 | 0.87 | Repressed | 1.05E-15 |
| Type I interferon signaling pathway | 48 | 2.17 | Induced | 1.55E-13 |
| Response to type I interferon | 52 | 2.06 | Induced | 1.55E-13 |
| Epidermal cell differentiation | 98 | 1.36 | Repressed | 5.09E-13 |
| rRNA metabolic process | 100 | 0.79 | Repressed | 5.09E-13 |
| Epidermis development | 137 | 1.11 | Repressed | 1.63E-12 |
| Keratinization | 66 | 1.71 | Repressed | 5.92E-12 |
| Keratinocyte differentiation | 87 | 1.36 | Repressed | 8.12E-11 |
| Humoral immune response | 84 | 1.35 | Induced | 1.02E-10 |
| Response to interferon-gamma | 95 | 1.45 | Induced | 1.14E-09 |
| Cornification | 59 | 1.60 | Repressed | 4.15E-09 |

The enhanced pro-inflammatory and pro-EMT response to infection we observe in HCECs relative to HVEs could be explained by altered basal expression of genes associated with these processes. Thus, we next compared the transcriptome of mock-infected control HCECs to that of mock-infected HVEs, identifying 4720 genes differentially expressed in an HCEC-intrinsic fashion (Figure 1B). Functional characterization of this gene set via STRING revealed a similar pattern of enriched terms as in the comparison of *Chlamydia*-infected cells, with relative induction of innate immunity as well as repression of keratinization and epidermal differentiation (Table 2). Expression of specific interleukins and CXCL-family cytokines followed a similar trend, with infected HCECs exhibiting strikingly increased expression of *CXCL10* (FC: 480.76), *IL6* (FC: 1423.03) and *IL8* (FC: 186.51), as well as modest induction of *IL1B* (FC: 3.73) and *IL24* (FC: 6.52) (Supplemental Table 1). However, expression of other interleukins, such as *IL7*, *IL15* and *IL20*, differed only between *Chlamydia*-infected HCECs and HVEs (Figure 1C). Importantly, both IL-15 and IL-20 have been implicated in fibrosis^35, 36^, suggesting that *Chlamydia*-infected endocervical epithelial cells may produce a wider portfolio of fibromodulatory cytokines relative to infection of other cell types. In contrast cytokeratin repression in HCECs was far more consistent between uninfected and *Chlamydia*-infected transcriptomes, as was relative induction of *SNAI1* and *TWIST1* (Figure 1C). These data suggest HCEC-specific expression of EMT-associated genes may be the product of an intrinsic predisposition towards EMT induction, rather than an altered response to infection.

**Table 2:** GO Biological Process terms significantly enriched (FDR ≤ 0.05) in comparison of steady-state expression in primary human endocervical epithelial cells (HCECs) and primary human vaginal epithelial cells (HVEs) via bulk RNA-sequencing. Induction/repression of each term is relative to HVE expression.

| <u>GO-BP Term Description</u> | <u>Genes Mapped</u> | <u>Enrichment Score</u> | <u>Modulation</u> | <u>False Discovery Rate</u> |
| --- | --- | --- | --- | --- |
| Innate immune response | 196 | 1.06082 | Induced | 1.04E-10 |
| Humoral immune response | 59 | 1.65343 | Induced | 1.07E-07 |
| Epidermal cell differentiation | 79 | 1.15647 | Repressed | 3.85E-07 |
| Epidermis development | 113 | 0.966933 | Repressed | 2.39E-06 |
| Epithelial cell differentiation | 191 | 0.58554 | Repressed | 1.42E-05 |
| Keratinization | 57 | 1.30416 | Repressed | 1.42E-05 |
| Cornification | 53 | 1.29258 | Repressed | 1.42E-05 |
| Response to bacterium | 168 | 0.767625 | Induced | 1.66E-05 |
| Cytokine-mediated signaling pathway | 200 | 0.872896 | Induced | 1.66E-05 |
| Regulation of defense response | 184 | 0.680786 | Induced | 1.66E-05 |
| Inflammatory response | 151 | 0.82777 | Induced | 2.73E-05 |
| Keratinocyte differentiation | 67 | 1.14483 | Repressed | 4.06E-05 |

### Cervical epithelial cells do not exhibit collagen or EMT induction during midcycle infection by *C. trachomatis*

While the effect of *C. trachomatis* infection on gene expression of host cervical epithelial cells has been extensively reported ^20, 21, 37^, prior study has focused on HPV-transformed cell lines (e.g. HeLa, HEp-2). Given the potential for HPV transformation to dysregulate gene expression^38, 39^, we next defined a more physiologically relevant host transcriptome by assessing differential expression of *Chlamydia*-infected primary HCECs. Relative to mock-infected control HCECs, infection with *C. trachomatis* serovar L2 induced differential expression (FDRP ≤ 0.05) of 8241 genes at 24 hpi (Supplemental Table 1), In accordance with prior reports describing the extensive pro-inflammatory response to infection, functional characterization of this gene set via STRING exhibited enrichment of immune and inflammation-associated GO-BP terms, including both positive and negative regulation of type I/II interferon signaling (Table 3). Critically, this included infection-dependent induction of specific interleukins (e.g. *IL17*, *IL15*, *IL20*) previously noted to exhibit HCEC-specific expression (Figure 2A).

**Figure 2.**
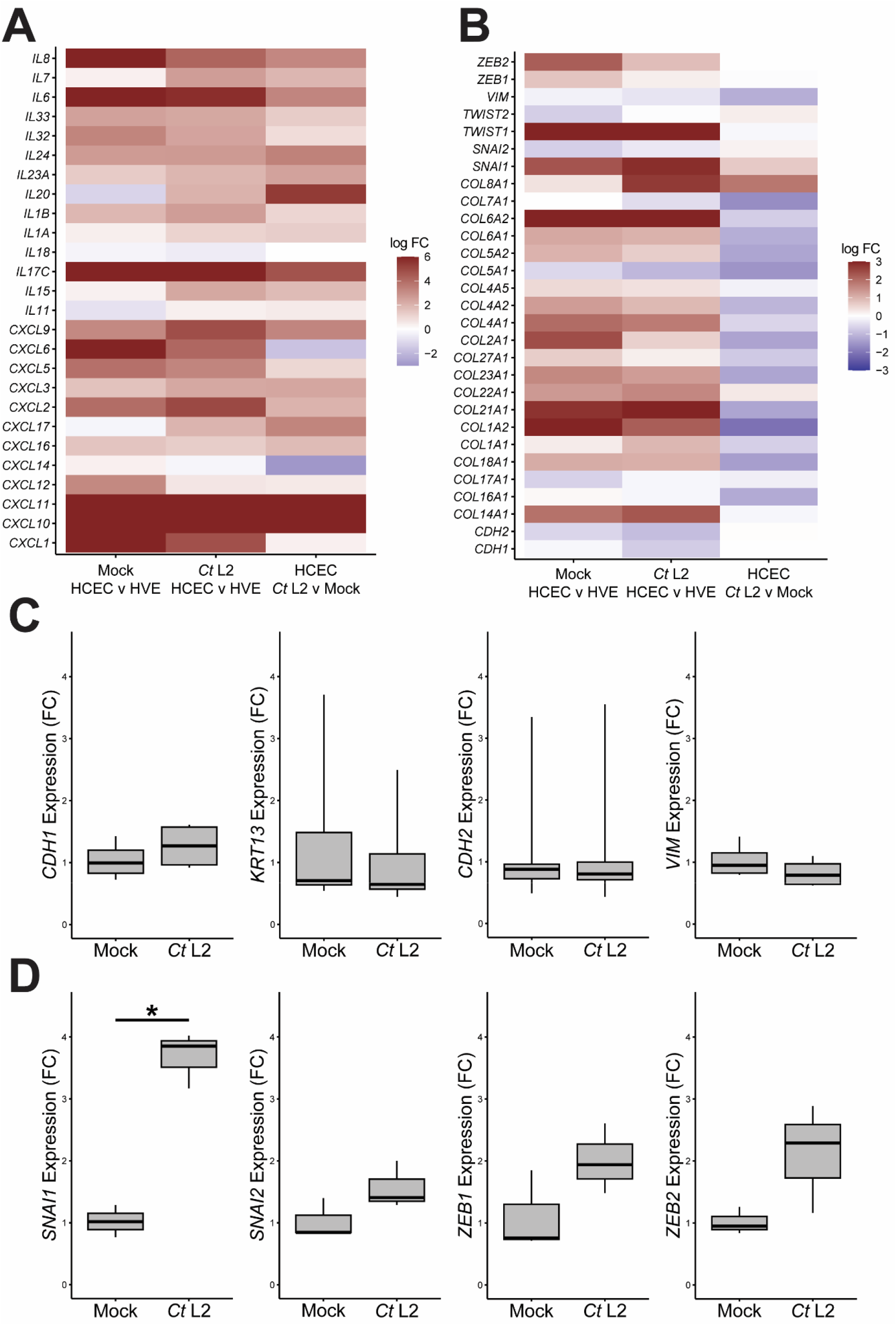
Cervical epithelial cells do not exhibit collagen or EMT induction during midcycle infection by *C. trachomatis*. **(A, B)** Heatmap of cytokine **(A)** and collagen/EMT-associated **(B)** gene expression in mock-infected HCECs relative to equivalent HVEs (left columns), *Ct* L2-infected HCECs relative to equivalent HVEs (center columns), and *Ct* L2-infected HCECs relative to mock-infected HCECs (right columns). Only genes differentially expressed (FDRP ≤ 0.05) in at least one comparison are shown. **(C, D)** Expression of epithelial/mesenchymal differentiation markers **(C)** and EMT-associated transcription factors **(D)** at 24 hpi in mock- and *Ct* L2-infected End1/E6E7 immortalized endocervical epithelial cells, as measured by RT-qPCR. n = 3 biological replicates; fold changes are relative to mean expression of the mock-infected and untreated control. Whiskers: minimum to maximum; asterisks: p-values ≤ 0.05, using pairwise Student’s t-tests and Bonferroni’s correction for multiple comparisons.

**Table 3:**
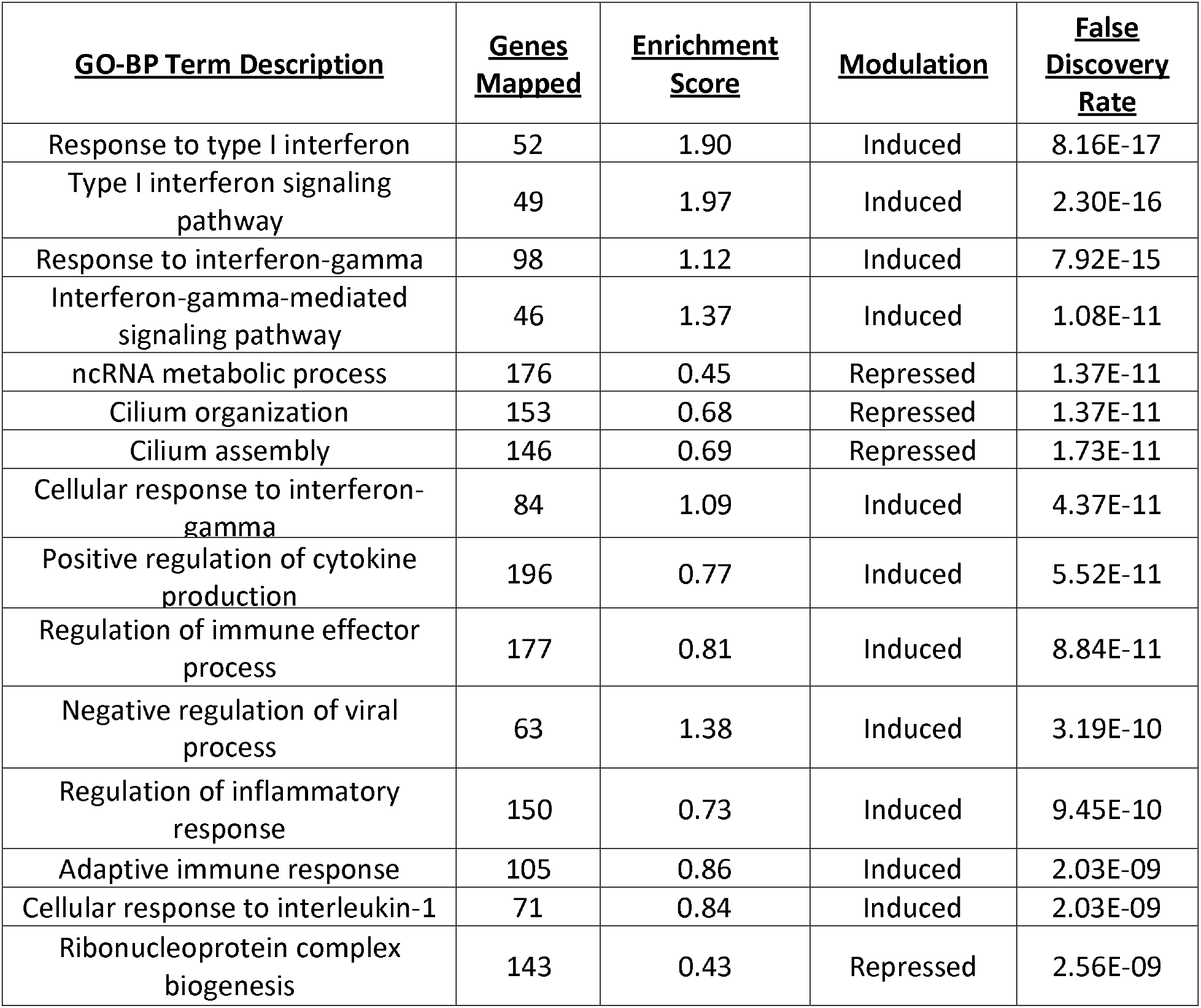
GO Biological Process terms significantly enriched (FDR ≤ 0.05) in comparison of *C. trachomatis* serovar L2-infected HCECs to mock-infected control HCECs via bulk RNA-sequencing. Induction/repression of each term is relative to the mock infection.

Intriguingly, the infection-associated transcriptome exhibited repression of extracellular matrix (ECM) components, with GO-CP (gene ontology, cellular component) terms associated with collagens and the ECM exhibiting modest enrichment amongst the downregulated gene set (Table 5). Indeed, infected HCECs exhibited reduced expression of fibrillar (e.g. *COL1A1*, *COL5A1*), fibril-associated (e.g. *COL16A1*, *COL21A1*), and network-forming (*COL4A1*) collagens relative to mock-infected HCECs – despite collagen expression of infected HCECs generally exceeding that of infected HVEs (Figure 2B). The mesenchymal marker vimentin/*VIM* was repressed by infection (FC: −2.22), whereas common markers of EMT (e.g. *CDH1*/2, *SNAI1*/*2*, *ZEB1*/*2*) were not differentially expressed (Figure 2B), suggesting that infection does not induce EMT in endocervical epithelial cells at 24 hpi. To confirm this hypothesis, we next examined expression of EMT-associated markers at this time point during *C. trachomatis* serovar L2 infection of immortalized endocervical epithelial cells (End1/E6E7), which have previously been demonstrated to undergo EMT in a chronic/persistent model of *C. trachomatis* infection ^40^. Critically, infected End1s did not exhibit induction of mesenchymal differentiation markers (e.g. CDH2, VIM) or repression of epithelial differentiation markers (e.g. CDH1, KRT13) (Figure 2C). In similar fashion, transcription factors associated with EMT induction did not exhibit induction at 24 hpi, with the notable exception of SNAI1/SNAIL (Figure 2D). Collectively, these data imply that chlamydial induction of EMT in cervical epithelial cells does not occur by 24 hpi, despite an apparent predisposition towards mesenchymal differentiation relative to vaginal epithelial cells. Thus, pro-fibrotic gene expression observed at this time point likely occurs via an EMT-independent mechanism.

### Induction of the pro-fibrotic transcriptional cofactor YAP is specific to infection of cervical epithelial cells

To identify alternative mechanisms of pro-fibrotic gene expression potentially modulated by *Chlamydia*, we next defined a common signature of pro-fibrotic genes exhibiting distinct expression in the upper genital tract as well as differential expression during *C. trachomatis* infection. Importantly, only a subset of genes differentially expressed in an HCEC-intrinsic fashion (mock HCEC vs mock HVE expression) exhibited comparable induction/repression in an infection-mediated fashion (mock HCEC vs *Ct* L2-infected HCEC expression). Upon comparing the HCEC-intrinsic (4720 genes) and infection-mediated (8241 genes) differentially expressed gene sets, only 2391 genes were common to both (Figure 3A). Of these, 1158 (49.4%) were equivalently induced or repressed in both comparisons, suggesting their likely relevance to infection-associated fibrosis of the upper genital tract (Figure 3A). More than half (629 genes, or 54.3%) of this gene set mapped to the Comparative Toxicogenomics Database of pro-fibrotic genes (Figure 3B), including numerous pro-fibrotic signal factors (e.g. *IL6*, *IL24, TGFA*) and ECM-remodeling enzymes (e.g. *ADAMTS9*, *MMP13*, *TIMP2/3*) (Supplemental Table 2). Critically, expression of this gene set in infected HCECs significantly correlated with our previously reported bulk RNA-sequencing of *Ct* serovar L2-infected End1/E6E7 immortalized endocervical epithelial cells (Figure 3B). These data suggest that modulation of this putatively fibrotic gene set is conserved between *in vitro* models of chlamydial pathogenesis in the upper genital tract, additionally recommending use of End1/E6E7 cells for subsequent investigation of the mechanisms underlying infection-associated pro-fibrotic gene expression.

**Figure 3.**
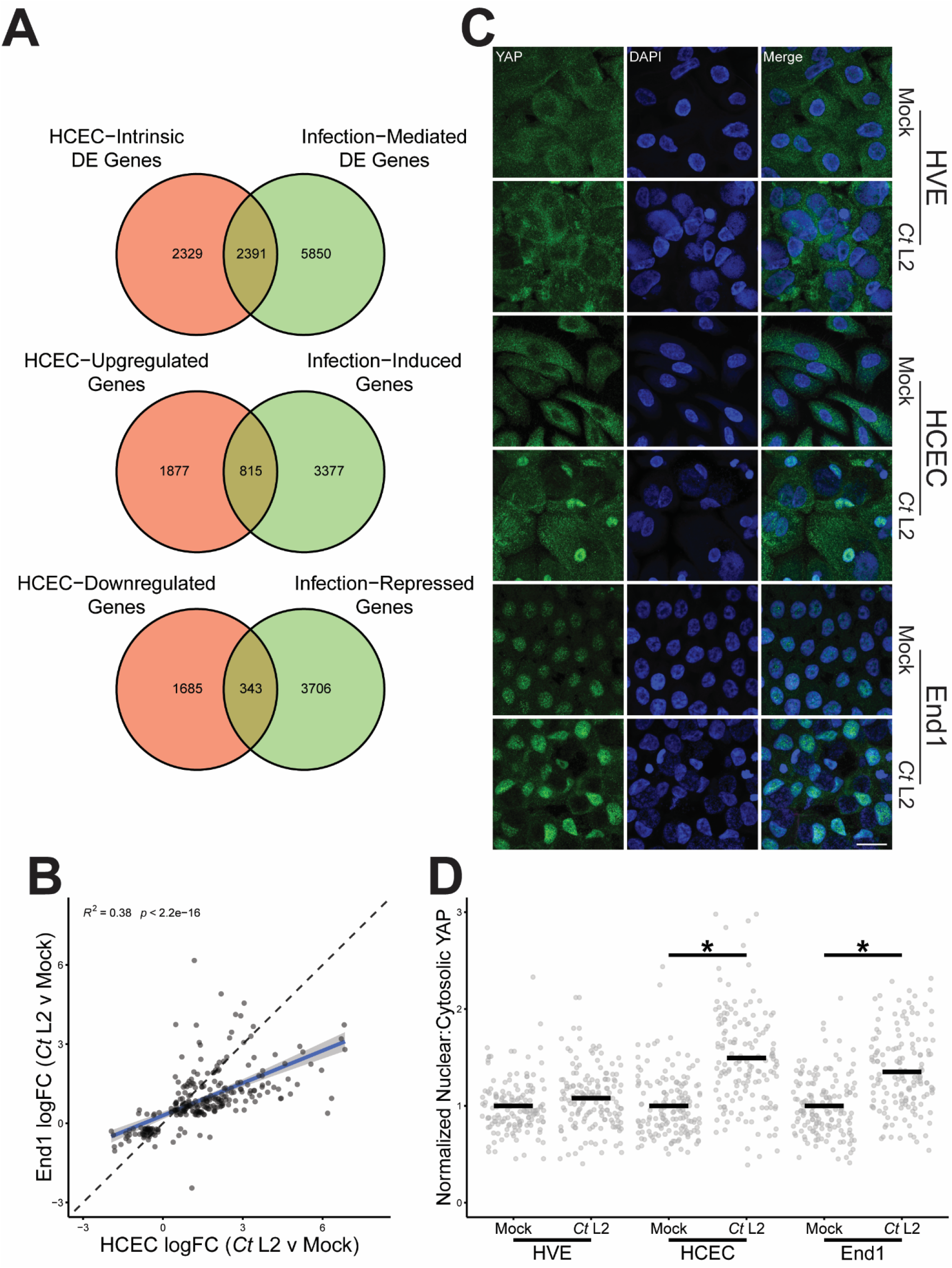
Induction of the pro-fibrotic transcriptional cofactor YAP is specific to infection of cervical epithelial cells. **(A)** Venn diagrams of genes differentially expressed (top, FDRP ≤ 0.05), induced (middle, FDRP ≤ 0.05, log_2_FC > 0) or repressed (bottom, FDRP ≤ 0.05, log_2_FC < 0) in an HCEC-intrinsic (red, mock-infected HCECs v mock-infected HVEs) or infection-mediated fashion (green, *Ct* L2-infected HCECs v mock-infected HCECs). **(B)** Scatter plot of gene expression *Ct* L2 infection of HCECs (x-axis) and End1s (y-axis) at 24 hpi, of the subset of genes equivalently modulated by steady-state HCEC expression relative to HVEs (see also Supplementary Table 2). All fold changes are relative to each cell type’s respective mock-infected control; blue line: linear regression model of correlation; grey shading: 95% confidence interval. R^2^ and p-values calculated using Pearson’s correlation. **(C)** Representative micrographs of YAP (green) translocation into the nuclei (blue) of confluent mock- and *Ct* L2-infected HVEs, HCECs, and End1s at 24 hpi. Scale bar: 20 μm. **(D)** Quantification of YAP nuclear translocation in **(C)** as a ratio of nuclear to cytosolic YAP fluorescence normalized to each cell type’s respective mock-infected control. n = 3 biological replicates, 50 cells measured per sample. Black bars: group means; asterisks: p-values ≤ 0.05, using pairwise Wilcoxon rank sum tests and Bonferroni’s correction for multiple comparisons.

One potential means by which infection may facilitate host expression of pro-fibrotic gene expression is via modulation of host transcription factors. Indeed, past work by our laboratory and others have defined mechanisms by which *C. trachomatis* and related species manipulate host transcription factor activity ^12, 41, 42^. Thus, we next performed transcription factor enrichment analysis of this putatively fibrotic gene set via ChEA3 (ChIP-X Enrichment Analysis 3) ^43^, identifying 56 transcription factors exhibiting significant enrichment (Supplemental Table 3). This included known mediators of pro-fibrotic and pro-inflammatory signaling, such as RELA, IRF1/8, SMAD3, and EGR1. Strikingly, nearly half (25) of the enriched transcription factors are known regulatory targets of YAP (Table 4), a pro-fibrotic transcriptional cofactor of which we recently reported *Chlamydia*-directed induction ^12^. Given the potential for YAP activation to drive expression of pro-fibrotic gene expression specific to the upper genital tract, we next assessed the degree to which *C. trachomatis* infection facilitates YAP nuclear translocation in vaginal epithelial cells. After infecting confluent monolayers of each cell type with *C. trachomatis* serovar L2, YAP nuclear translocation was quantified as a ratio of nuclear to cytosolic immunofluorescence. Critically, infected HVEs did not exhibit YAP nuclear translocation relative to mock-infected control monolayers, in stark contrast to the significant increase in nuclear YAP observed during infection of either primary (HCEC) or immortalized (End1) endocervical epithelial cells (Figures 3C, 3D). Collectively, these data suggest that chlamydial YAP activation may mediate pro-fibrotic gene expression specific to infection of the upper genital tract.

**Table 4:** YAP-associated transcription factors exhibiting significant enrichment (FDR ≤ 0.05) in the set of genes equivalently modulated both by *Ct* L2 infection of HCECs and steady-state HCEC expression relative to HVEs.

| <u>Transcription Factor</u> | <u>Genes Mapped</u> | <u>Set Length</u> | <u>FDR</u> | <u>YAP Regulatory Relationship</u> |
| --- | --- | --- | --- | --- |
| SOX2 | 236 | 2308 | 8.68E-21 | Expression induced by YAP/Oct4 <sup>76</sup> |
| ESR2 | 50 | 394 | 4.81E-08 | Expression repressed by YAP/NuRD <sup>77</sup> |
| BACH1 | 114 | 1298 | 1.96E-07 | Coactivated by YAP <sup>78</sup> |
| STAT3 | 82 | 914 | 5.90E-06 | Coactivated by YAP <sup>79</sup> |
| JUN | 138 | 1830 | 2.60E-05 | Coactivated by YAP <sup>79</sup> |
| SMAD3 | 129 | 1759 | 1.56E-04 | Coactivated by YAP <sup>66</sup> |
| JUND | 128 | 1831 | 8.94E-04 | Coactivated by YAP <sup>79</sup> |
| SALL4 | 62 | 775 | 0.00155 | Expression induced by YAP/TEAD2 <sup>80</sup> |
| MYB | 64 | 837 | 0.0036 | Coactivated by YAP <sup>81,82</sup> |
| EGR1 | 49 | 605 | 0.00438 | Coactivated by YAP <sup>83</sup> |
| ATF3 | 130 | 1971 | 0.00541 | Expression induced by YAP/TAZ <sup>84</sup> |
| FOXA2 | 178 | 2814 | 0.00541 | Enhancer binding altered by YAP <sup>80</sup> |
| SOX9 | 89 | 1270 | 0.00541 | Induced/activated by YAP/TEAD <sup>85</sup> |
| NANOG | 157 | 2452 | 0.00582 | Expression induced by YAP/TEAD2 <sup>86</sup> |
| WT1 | 99 | 1450 | 0.00612 | Represses CDH1 with YAP <sup>87</sup> |
| CEBPB | 90 | 1300 | 0.0064 | Coactivated by YAP <sup>79</sup> |
| RXRA | 106 | 1584 | 0.00738 | Coactivated by YAP <sup>88</sup> |
| TP53 | 69 | 981 | 0.0122 | Expression induced by YAP <sup>89</sup> |
| CDX2 | 37 | 461 | 0.0141 | Expression induced by YAP/TEAD4 <sup>90</sup> |
| MITF | 300 | 5140 | 0.0141 | Expression induced by YAP/PAX3 <sup>91</sup> |
| SMAD4 | 31 | 371 | 0.0158 | YAP-dependent nuclear retention <sup>92</sup> |
| RARG | 29 | 342 | 0.0165 | Coactivated by YAP <sup>88</sup> |
| KLF5 | 113 | 1803 | 0.0271 | Induced/stabilized by YAP <sup>93</sup> |
| FOXA1 | 113 | 1842 | 0.0421 | Bound by YAP/TFCP2 complex <sup>94</sup> |
| CEBPA | 36 | 493 | 0.0461 | Coactivated by YAP <sup>79</sup> |

### Coculture with infected endocervical epithelial cells alters the putative myofibroblast transcriptome

Given our data indicating that infected endocervical epithelial cells exhibit induction of pro-fibrotic signal factors, one potential mechanism through which *C. trachomatis* may induce fibrosis is via signaling to stromal fibroblasts. Indeed, differentiation of tissue-resident fibroblasts into ECM-remodeling myofibroblasts has been consistently associated with other forms of fibrotic disease^26^; however, the contribution of fibroblasts to Chlamydia-associated scarring has not been assessed. Importantly, the lack of clinical evidence of *Chlamydia*-infected fibroblasts *in vivo* necessitates a model of infection wherein uninfected fibroblasts are cocultured with infected endocervical epithelial cells. Past study of the kinetics of *C. trachomatis* infection epithelial cells indicates that pathogen-directed uptake of the bacterium is rapid, with a majority of invading bacteria internalized as soon as 10 minutes post-infection ^44, 45^. Critically, this event requires interaction of *Chlamydia* with the host cell plasma membrane in two distinct stages: transient adhesion relying on electrostatic interactions with membrane glycosaminoglycans ^46, 47^, followed by irreversible attachment due to binding with host cell receptors and/or engagement of the pathogen’s type III secretion system^48^. Thus, we infected immortalized End1s seeded at 35% of confluence with the genetically tractable *C. trachomatis* serovar L2, adding immortalized KCO2-44D uterine fibroblasts at one hour post-infection (hpi). After 23 hours of incubation, cocultures prepared via this method exhibited a consistent morphology characterized by islands of epithelial cells interspersed with fibroblasts, of which the latter were readily distinguished by immunofluorescence staining of actin stress fibers (Figure 4A). Importantly, fibroblasts did not exhibit visible chlamydial inclusions containing DAPI-positive organisms, in contrast to the readily apparent inclusions of infected epithelial cells (Figure 4A).

**Figure 4.**
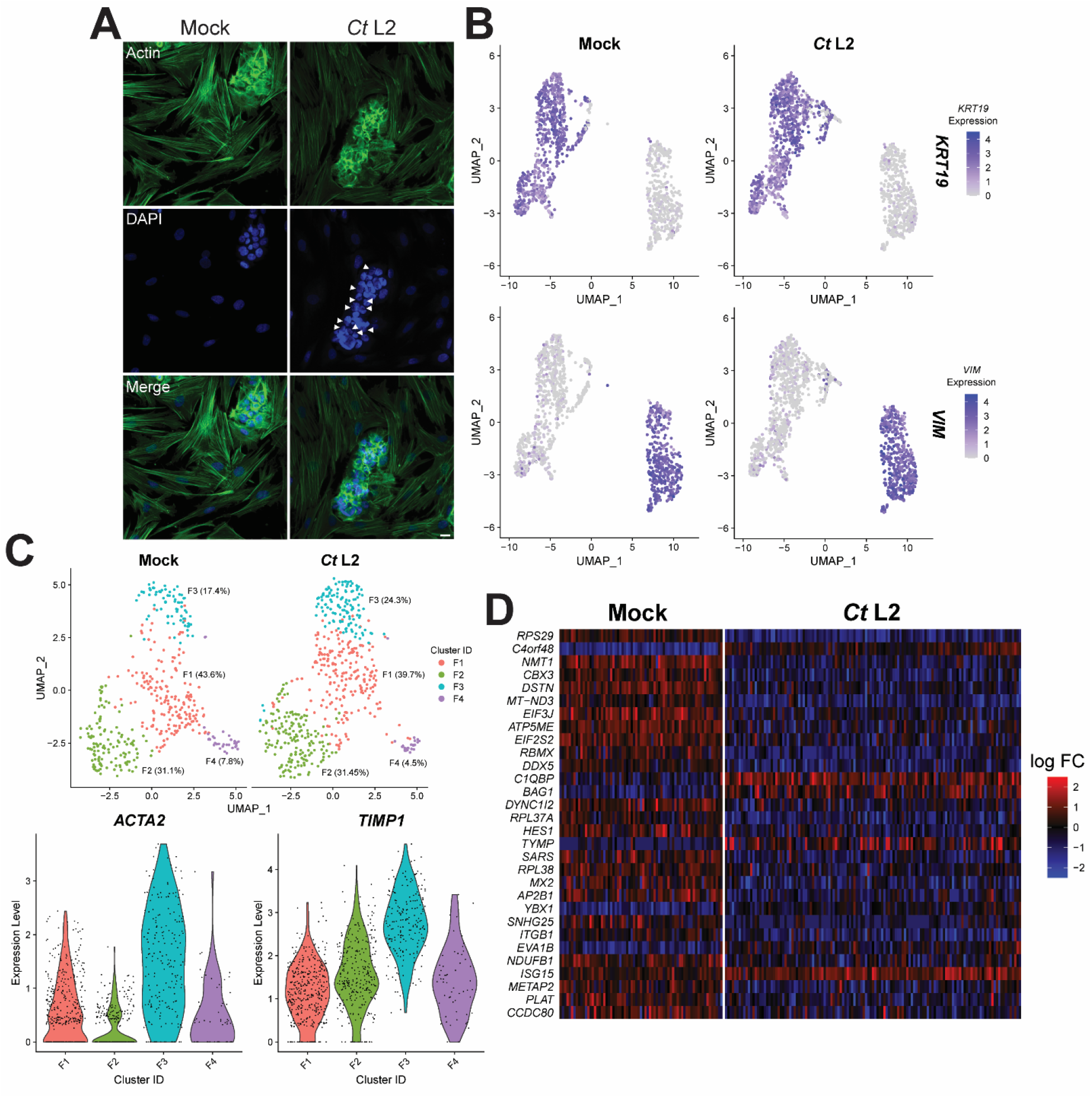
Coculture with infected endocervical epithelial cells alters the putative myofibroblast transcriptome. **(A)** Representative micrographs of mock-(left) or C. trachomatis serovar L2-infected (right) End1/E6E7 endocervical epithelial cells cocultured with KCO2 uterine fibroblasts for 23 h (starting at 1 hpi); green: actin expression (phalloidin), blue: DAPI, arrowheads: chlamydial inclusions, scale bar: 20 μm. **(B)** Uniform manifold approximation and projection (UMAP) feature plot of *KRT19* (top) and *VIM* (bottom) expression in same-well coculture of mock- or *Ct* serovar L2-infected End1 endocervical epithelial cells with KCO2 fibroblasts for 23 h (starting at 1 hpi). n = 2 biological replicates, 1000-cell libraries per sample, approx. 5-7 x 10^4^ reads/cell. **(C)** (top) UMAP plot of fibroblast (*VIM*+) subclustering in same-well coculture of mock- or *Ct* serovar L2-infected End1 endocervical epithelial cells with KCO2 fibroblasts. Colors: cluster identity, all percentages relative to total fibroblast population of the respective mock-/*Ct* L2-infected coculture. (bottom) Violin plot of fibroblast (*VIM*+) per-cluster expression of α-SMA/*ACTA2* (left) and *TIMP1* (right) in same-well coculture of mock- or *Ct* serovar L2-infected End1 endocervical epithelial cells with KCO2 fibroblasts. Colors: cluster identity, dots: per-cell expression values. **(D)** Per-cell expression heatmap of genes differentially expressed (p ≤ 0.05) by cluster F3 fibroblasts of *Ct* serovar L2-infected cocultures (right), relative to cluster F3 fibroblasts of mock-infected cocultures (left).

To both confirm the absence of Chlamydia-infected fibroblasts and examine the transcriptome of uninfected fibroblasts, we then harvested cocultures at 24 hpi via trypsinization for library preparation and single-cell RNA-sequencing. End1/KCO2 cell populations were readily distinguished on the basis of epithelial and mesenchymal marker expression, with End1s exhibiting heightened expression of cytokeratin 19 (KRT19) and KCO2s exhibiting heightened expression of vimentin (VIM) (Figure 4B). To identify Chlamydia-infected cells, sequencing results were aligned to the hg18 reference genome appended with C. trachomatis reference sequences for highly expressed, polyadenylated RNA transcripts most likely to be captured by library preparation and sequencing, including the chlamydial outer membrane protein ompA, the polymorphic membrane protein pmpC, the nucleoside phosphate kinase ndk, and the pgp8 antisense sRNA-2. Consistently, expression of chlamydial reference sequences identified an epithelial cell cluster distinct from the mesenchymal population, and was not detectable in uninfected cocultures (Supplemental Figure 1). Taken together, these data demonstrate the viability of this approach as means to render cocultured fibroblasts inaccessible to Chlamydia.

To determine if coculture with infected epithelial cells altered the transcriptome of “bystander” fibroblasts, we performed clustering analysis on the mesenchymal cell population via Seurat. We identified four mesenchymal subclusters distinguished by marker gene expression, designated clusters F1-F4 (Figure 4C, top), with the mesenchymal population of Chlamydia-infected cocultures exhibiting a modest increase in the proportion of cluster F3 relative to a mock-infected control (24.3% and 17.4%, respectively). Given that cluster F3 also exhibited enhanced expression of the myofibroblast differentiation marker smooth muscle actin (α-SMA/ACTA2), as well as tissue inhibitor of matrix metallopeptidases 1 (TIMP1), this result suggests that coculture with infected-epithelial cells may enhance myofibroblast differentiation (Figure 4C, bottom). Further comparison of cluster F3 gene expression in Chlamydia- and mock-infected cocultures revealed differential expression of a variety of genes associated with fibrosis (Figure 4D). For example, cluster F3 fibroblasts from Chlamydia-infected cocultures exhibited significant repression of HES1, a bHLH-family transcription factor known to inhibit the regulator of myofibroblast differentiation MyoD^49, 50^. Coculture with infected epithelial cells additionally inhibited expression of chromobox protein homolog 3 (CBX3), which was recently shown to inhibit proliferation and collagen production of vascular smooth muscle cells ^51^. “Bystander” cluster F3 fibroblasts also exhibited increased expression of pro-inflammatory genes, including C1QBP and ISG15, suggesting that cocultured fibroblasts may respond to infection-associated induction of pro-inflammatory signaling by host endocervical epithelial cells. Collectively, these data imply that infected epithelial cells may signal to cocultured fibroblasts, thereby stimulating pro-fibrotic gene expression.

### *Chlamydia*-infected endocervical epithelial cells induce fibroblast collagen I expression in a YAP-dependent fashion

A key functional consequence of myofibroblast differentiation is the capacity to remodel the extracellular matrix via the production of collagens ^52, 53^. In healthy epithelial tissues, the ECM of the basement membrane is predominantly composed of type IV network-forming collagen, laminins, and the collagen/laminin crosslinking protein nidogen ^54, 55^. In contrast, fibrotic tissues exhibit heightened proportionality of fibrillar collagens (e.g. types I, II, III, V) in the basement membrane, which has been attributed to deposition by myofibroblasts in idiopathic pulmonary fibrosis and other forms of scarring disease ^52, 54, 56, 57^. In similar fashion, it has been shown that *Chlamydia*-infected epithelial cells exhibit increased production of fibrillar collagen due to chlamydial induction of EMT and consequent host cell transdifferentiation into a myofibroblast-like phenotype^17^.

In light of these results, one potential consequence of the altered fibroblast transcriptome in *Chlamydia*-infected cocultures may be increased production of fibrillar collagens. Thus, we prepared an alternative *in vitro* model of chlamydial infection, wherein End1 epithelial cells were seeded/infected on the semipermeable membrane of a transwell insert, washed with heparan sulfate at 1 hpi, then transferred to a plate containing seeded KCO2 fibroblasts (Figure 5A). This approach provided a means to perform population-level measurement on cocultured fibroblasts, given that the 0.4 μm pore size of the transwell membrane would necessarily restrict passage of cells, while still allowing for cell-cell communication via the action of diffusible signal factors. Critically, KCO2s cocultured with *Ct* serovar L2-infected End1s in transwells for 23 hours (24 hpi) exhibited a significant increase in production of type I collagen relative to an uninfected control, as measured by immunofluorescence (Figure 5B, 5C). Induction of collagen I expression occurred in a fashion consistent with mock-infected cocultures treated with TGF-β1 (a known inducer of myofibroblast differentiation and collagen deposition) ^58–60^.

**Figure 5.**
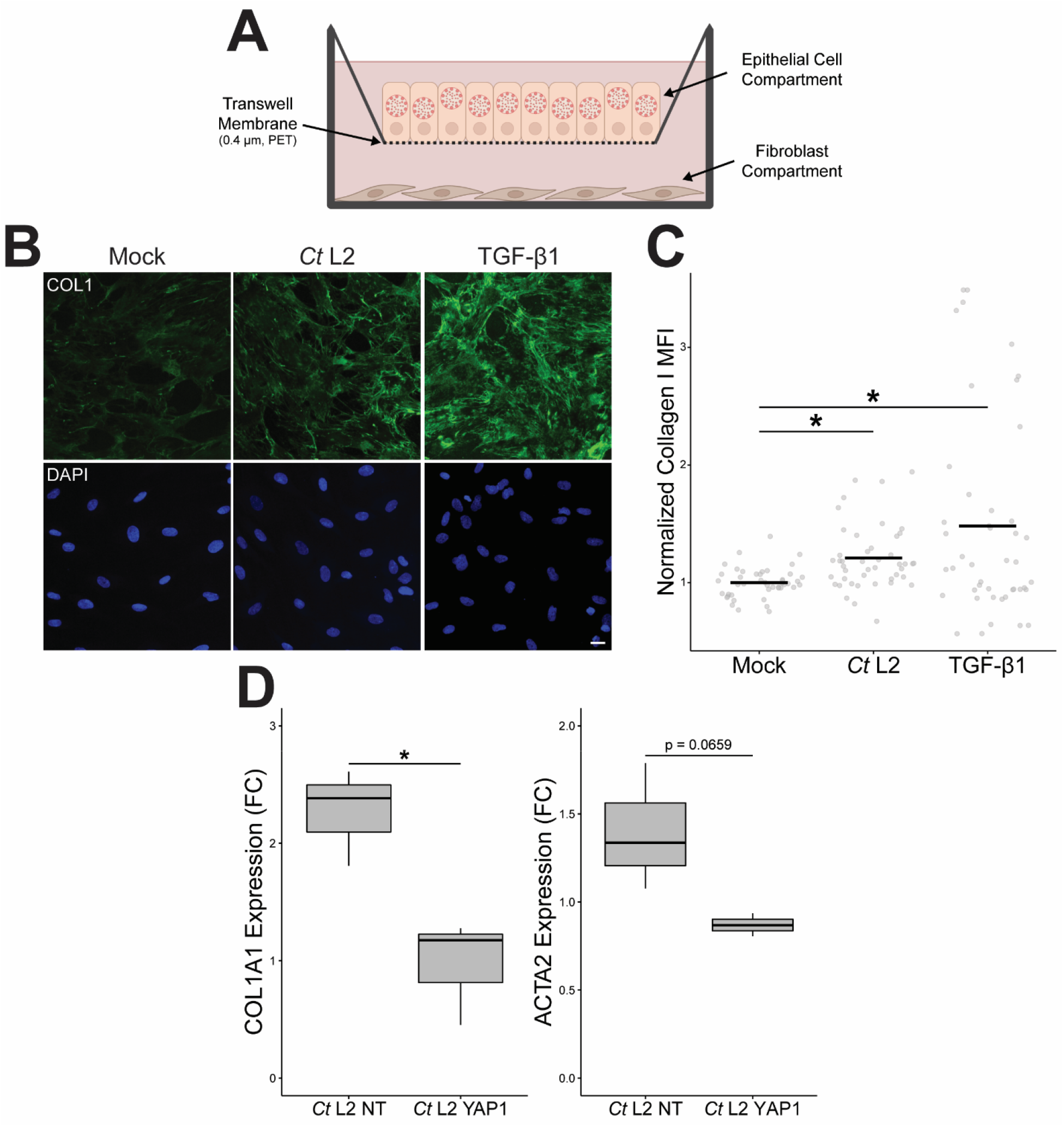
*Chlamydia*-infected endocervical epithelial cells induce fibroblast expression of collagen I in a YAP-dependent fashion. **(A)** Schematic representation of the transwell-mediated approach of coculturing *C. trachomatis*-infected endocervical epithelial cells (top compartment) with uninfected uterine fibroblasts (bottom compartment). **(B)** Representative micrographs of collagen I (green) produced by KCO2 fibroblasts cocultured with mock-infected, *Ct* L2-infected, or TGF-β1-treated (20 ng/mL) End1 cells for 24 h, starting at 1 hpi. Scale bar: 20 μm. **(C)** Quantification of mean fluorescence intensity (MFI) of collagen I in (B). n = 3 biological replicates, 5 fields measured per sample. Black bars: group means; asterisks: p-values ≤ 0.05, using pairwise Wilcoxon rank sum tests and Bonferroni’s correction for multiple comparisons. **(D)** *COL1A1* and *ACTA2* expression as measured by RT-qPCR in KCO2 fibroblasts cocultured for 24 h with siRNA-transfected (10 nM of YAP1-targeting or non-targeting siRNA, for 24 h prior to infection),*Ct* L2-infected End1 cells (coculture starting at 1 hpi). n = 3 biological replicates; fold changes are relative to mean expression of the equivalent mock-infected control. Whiskers: minimum to maximum; asterisks: p-values ≤ 0.05, using Student’s t-tests and Bonferroni’s correction for multiple comparisons.

We have previously demonstrated that *C. trachomatis* infection of endocervical epithelial cells induces expression of the fibroblast-activating signal factors *CTGF* and *INHBA*, in a fashion dependent upon pathogen-directed induction of the host transcriptional cofactor YAP^12^. Further, here we observe infection-mediated YAP activation specific to cervical epithelial cells, suggesting a role for chlamydial YAP activation in the fibrotic pathology associated with chlamydial infection of the upper genital tract.Thus, to assess the role of chlamydial YAP activation in stimulation of fibroblast collagen production, we next opted to knock down YAP expression in epithelial cells via siRNA at 24 hours prior to infection. To confirm our previous results using an alternate method, fibroblast collagen expression after 23 hours of coculture (24 hpi) was assessed via RT-qPCR. Knockdown of YAP in *Chlamydia*-infected End1s was approximately 40% efficient, as measured by Western blotting of total YAP protein levels – in accordance with other reports of siRNA-mediated knockdown efficiency in End1s ^12, 61^ (Supplemental Figure 2). Fibroblast expression of collagen I was significantly increased in infected cocultures transfected with non-targeting control siRNA, consistent with our prior observations via immunofluorescence. However, transfection of infected End1s with YAP-targeting siRNA significantly attenuated this phenotype, with cocultured fibroblasts exhibiting collagen I (*COL1A1*) expression indistinguishable from mock-infected cocultures (Figure 5D). Intriguingly, expression of the myofibroblast differentiation marker α-SMA (*ACTA2*) exhibited a similar, albeit statistically insignificant trend (Figure 5D). Taken together, these data confirm that *Chlamydia*-infected epithelial cells stimulate fibroblast production of collagen, as well as implicate chlamydial induction of host cell YAP activity in this effect.

## Discussion

To elucidate the mechanisms underlying *Chlamydia*-mediated fibrosis of the upper genital tract, we have presented comparative analysis of both steady-state and infection-associated gene transcription in primary human cervical epithelial cells (HCECs) and primary human vaginal epithelial cells (HVEs). Both mock- and *Chlamydia*-infected HCECs exhibited enhanced expression of pro-inflammatory and pro-fibrotic cytokines (e.g. *IL6*, *IL8*), induction of the pro-EMT transcription factors SNAI1/SNAIL and TWIST1, and relative repression of multiple cytokeratins. However, expression of specific fibromodulatory interleukins (eg. *IL7*, *IL15*, *IL20*) was enhanced only in *Chlamydia*-infected HCECs, suggesting a cell type-specific response to infection. Transcription factor enrichment analysis of genes equivalently modulated by infection and HCEC-intrinsic expression relative to HVEs revealed a broad portfolio of known YAP regulatory targets. YAP nuclear translocation of YAP was enhanced during infection of HCECs, but not HVEs, suggesting a role for this transcriptional cofactor in mediating the HCEC-specific induction of pro-fibrotic signaling. To investigate this possibility, we examined the response of uninfected fibroblasts to coculture with *Chlamydia*-infected endocervical epithelial cells, observing a modest expansion of α-SMA/*TIMP1*-expressing putative myofibroblasts. Further iteration on this coculture approach via use of transwells enabled population-level measurement of cocultured fibroblast gene expression. We find that cocultured fibroblasts exhibit heightened expression of type I collagen – critically, this phenotype was sensitive to siRNA-mediated knockdown of YAP in infected endocervical epithelial cells. Taken together, our results present a novel mechanism by which *Chlamydia* mediates fibrotic tissue remodeling, wherein infection-mediated YAP activation in host endocervical epithelial cells promotes fibroblast collagen expression via the induction of pro-fibrotic signaling.

The apparent repression of collagen expression in infected HCECs is surprising, given past reports of chlamydial EMT leading to production of type I collagen in infected primary cells of the murine oviduct ^17^. Deposition of fibrillar collagen is a hallmark of pro-fibrotic remodeling of the ECM in other forms of scarring disease ^54–56^; in light of this, the role of *Chlamydia*-infected epithelial cells in collagen synthesis is critically understudied. These data suggest two possibilities: either previous reports of fibrillar collagen production by infected cells are unique to infection of murine oviduct epithelial cells, or infection inhibits collagen expression in host cells prior to induction of EMT. Importantly, prior study of chlamydial EMT induction indicates this phenotype occurs in late-cycle or persistent models of infection, with murine oviduct epithelial cells and End1 cells expressing mesenchymal markers at 72 hours or 7 days post-infection, respectively. Consequently, infection-mediated repression at 24 hpi may be a transient effect, superseded by pathogen-directed host transdifferentiation into a myofibroblast-like and collagen-producing phenotype. Indeed, prior reports indicating collagen expression is not enhanced in HeLa cells at 24 hpi would seem to support this hypothesis ^20^. Further study is required to elucidate the impact of chlamydial EMT induction on basement membrane architecture, and thereby determine the physiological relevance of this phenotype to the development of infection-associated fibrosis. Importantly, the infection-mediated induction of fibroblast collagen expression we observe may constitute a means by which *Chlamydia* may stimulate ECM deposition in an EMT-independent fashion.

Past study of *C. trachomatis* in murine models of infection has shown that the pathogen induces EMT in host epithelial cells of the murine oviduct, as well as host production of collagen I and TGF-β1 (a known inducer of myofibroblast differentiation) ^13, 16, 17^. However, fibrotic sequelae arising from *C. trachomatis* infection of the murine female genital tract has not been reported, suggesting that animal models of infection may not faithfully recapitulate the molecular events underlying *Chlamydia*-associated scarring in humans. In light of this, use of immortalized human cell lines in an *in vitro* model of infection thus has the potential to provide greater insight into how *Chlamydia*-infected cervical epithelial cells drive tissue-wide fibrotic pathologies. Here, we report that infected endocervical epithelial cells alter gene expression and induce collagen production in cocultured, uninfected fibroblasts – demonstrating a means by which *C. trachomatis* may coordinate pro-fibrotic activity of cell types beyond the simple columnar cervical epithelium the pathogen demonstrably infects *in vivo*.

The observation that collagen I expression in fibroblasts was sensitive to siRNA-mediated knockdown of YAP in epithelial cells is in accordance with our prior report of infection- and YAP-dependent expression of pro-fibrotic signal factors ^12^. However, it is unclear to what extent specific signaling factors are relevant to or dispensable for induction of this phenotype. Given consistent role in induction of fibroblast differentiation and ECM remodeling ^25, 62–64^, this signal factor constitutes a likely candidate for YAP-and infection-dependent induction of collagen synthesis in fibroblasts in cocultured fibroblasts. That being said, YAP has also been implicated in the expression of several other pro-fibrotic signaling factors. A recent report implicated YAP in inducing expression of the pro-fibrotic interleukin IL-6 in the context of endometrial cancer ^65^. YAP has also been shown to enhance TGF-β expression and signal transduction via binding to and enhancing the activity of SMAD2/3 ^66^. A recent report implicated YAP activity in expression of the myofibroblast-activating interleukin IL-33, with Cre/Lox-mediated YAP knockout significantly attenuating IL-33 expression and scarring in a murine model of post-infarction cardiac fibrosis ^67^. YAP has been additionally shown to modulate the Wnt/β-catenin signaling pathway, via enhancement of both β-catenin expression and transcriptional activity ^68^. Given the demonstrable role of this signaling pathway in fibrosis generally and regulation of TGF-β expression specifically^69, 70^, these data collectively demonstrate multiple, complementary means by which chlamydial induction of YAP may facilitate pro-fibrotic signaling. Ultimately, further study of the role of YAP must begin with clearly defining the YAP regulon in *Chlamydia*-infected endocervical epithelial cells. This may necessitate the development of a YAP-knockout cell line, given the relative inefficiency of siRNA-mediated knockdown we and others observe in End1/E6E7 cells ^61^.

It is important to note that chlamydial fibrosis is clinically associated with chronic or repeated infections, with the latter demonstrably increasing the risk of scarring infertility and/or ectopic pregnancy ^6^. Critically, the lack of a viable animal model of infection-associated scarring has heretofore precluded more longitudinal study of the pro-fibrotic effects of chronic or repeated *C. trachomatis* infection. Given our observation that *Chlamydia*-infected cells stimulate cocultured fibroblast expression of collagen, these data suggest intriguing possibilities: that infected epithelial cells sensitize affected tissues to other pro-fibrotic stimuli, or that repeated infections at the same site have an aggregate effect in stimulating scar-associated tissue remodeling. In a similar fashion, collagen deposition facilitated by infected epithelial cells may itself act as a stimulus of myofibroblast activation. Ultimately, the capacity of coculture-stimulated fibroblasts to return to a naïve, collagen-repressed state must be assessed, as must the potential for successive rounds of coculture to enhance fibroblast activation and collagen synthesis. Further refinement of the *in vitro* model of chlamydial pathogenesis presented here may provide critical insight into how the pro-fibrotic stimulus of infected epithelial cells may effectively persist after host cell death and pathogen clearance, thereby facilitating the tissue-level scarring associated with chronic or persistent infection.

## Materials and Methods

### Eukaryotic Cell Culture

Primary human cervical epithelial cells (HCECs, ATCC PCS-0480-011, Lot 80306190) were cultured at 37° C with 5% atmospheric CO_2_ in Cervical Epithelial Cell Basal Medium (CECBM, ATCC PCS-480-032) supplemented with all contents of a Cervical Epithelial Growth Kit (ATCC PCS-080-042), per manufacturer’s instructions. Primary human vaginal epithelial cells (HVEs, ATCC PCS-480-010, Lot 80924222) were cultured at 37° C with 5% atmospheric CO_2_ in Vaginal Epithelial Cell Basal Medium (VECBM, ATCC PCS-480-030) supplemented with all contents of a Vaginal Epithelial Cell Growth Kit (ATCC PCS-480-040), per manufacturer’s instructions. Human endocervical epithelial HPV-16 E6/E7-immortalized End1s (End1 E6/E7, ATCC CRL-2615) were cultured at 37° C with 5% atmospheric CO_2_ in Keratinocyte Serum-Free Medium (KSFM, Thermo Fisher Scientific) supplemented with human recombinant epidermal growth factor, bovine pituitary extract, 5 micrograms/mL gentamicin, and 0.4 mM CaCl_2_ (unless otherwise indicated). Human endometrial hTERT-immortalized fibroblasts (KCO2-44D hTERT, ATCC SC-6000) were cultured in Dulbecco’s modification of Eagle’s medium (DMEM, Thermo Fisher Scientific 11960069) supplemented with 10% Fetal Bovine Serum (FBS, Fisher Scientific NC0327704), 2 mM L-Glutamine (Thermo Fisher Scientific 25030081), and 5 ug/mL gentamicin. Both End1s and KCO2s were cultured at 37 C in 5% CO_2_, between passages 3-15. All cell lines were tested annually for mycoplasma contamination, using the ATCC PCR-based Mycoplasma Detection Kit (ATCC 30-1012K), per manufacturer’s instructions.

### Chlamydial Infections

*Chlamydia trachomatis* serovar L2 (434/Bu) was originally obtained from Dr. Ted Hackstadt (Rocky Mountain National Laboratory, NIAID). Chlamydial EBs were isolated from infected, cycloheximide-treated McCoy cells at 36-40 hours post-infection (hpi) and purified by density gradient centrifugation as previously described ^71^. For infection of 6-well plates (Greiner Bio-One 657160 and 662160), HCECs or HVEs seeded at 5000 cells/cm^2^ and grown to confluence (6-7 days) or End1s seeded at 125% of confluence and incubated for 16 h were washed with pre-warmed Hanks Buffered Saline Solution (HBSS) prior to inoculation with *Chlamydia*-containing CECBM, VECBM, or KSFM (respectively) at the indicated multiplicity of infection (MOI). To synchronize infection of cultured monolayers, tissue culture plates were centrifuged at 4° C and 500 rcf (Eppendorf 5810 R tabletop centrifuge, A-4-81 rotor) for 15 minutes, to permit transient electrostatic adherence of *C. trachomatis* to the host cell surface. Inoculum was then aspirated, and cells were washed with chilled HBSS to remove any non-adherent EBs. After aspiration of the wash, chlamydial invasion was initiated via the addition of CECBM, VECBM, or KSFM (HCECs, HVEs, or End1s respectively) pre-warmed to 37° C. Infected cultures were then incubated for 24 h at 37° C and 5% CO_2_ for subsequent harvesting of total RNA (see below).

For infection of End1/KCO2 same-well co-cultures, End1s were seeded at 35% of confluence on 6-well tissue culture plates (Greiner Bio-One 657160), incubated for 16 hours, then infected at an MOI of 5 as described above. Infected End1 cultures were returned to a tissue culture incubator until 1 hpi, at which time cultures were washed three times with pre-warmed HBSS containing 100 ug/mL heparin (Sigma-Aldrich H3393) to remove any transiently adhered, non-internalized EBs ^72^. After aspiration of the final wash, KCO2s in complete DMEM were seeded in each well, at 75% of confluence. Tissue culture plates were subsequently incubated until 24 hpi prior to harvest for single-cell RNA library preparation (see below).

For transwell coculture experiments, End1s seeded at 100% of confluence in 6-well-sized transwell inserts (Costar 3450, Fisher Scientific) and incubated for 16h (or 40h, in the case of experiments involving siRNA transfection) were infected at an MOI of 5 as above. At 1 hpi, cells were washed with HBSS containing 100 ug/mL heparin (Millipore-Sigma H3393) to remove transiently adhered chlamydial organisms. After replacement of wash media with DMEM, transwells were subsequently transferred to 6-well plates containing KCO2s seeded at 60% of confluence immediately after infection. Cocultures were then incubated for 24 h prior to fixation or RNA harvest (see below).

### Bulk RNA-Sequencing and Analysis

HVEs infected at an MOI of 2 as described above were harvested for RNA at 24 hpi using TRIzol (Thermo Fisher Scientific 15596026) and the DNA-free DNA removal kit (Thermo Fisher Scientific AM1906), according to manufacturers’ protocols. Total RNA samples were subsequently assayed for fragmentation using an Agilent Bioanalyzer 2100. Polyadenylated transcript enrichment and cDNA library preparation of intact samples was performed using the NuGEN Universal mRNA-Seq Library Preparation kit, and sequencing of cDNA libraries was performed using the NextSeq 550 system (Illumina). HCEC and End1 libraries were prepared as described previously ^12^.

Read alignment and downstream analysis was performed using CLC Genomics Workbench (Qiagen); each treatment group was comprised of libraries from three biological replicates, each with a minimum of 30 million reads (unstranded single read, mean length 150 bp), genes with an FDRP ≤ 0.05 were considered differentially expressed. Functional characterization of each differentially expressed gene set was performed using the STRING database of protein-protein association networks, using the ranked protein list functional enrichment analysis function ^31, 32^. GO/Wikipathway overrepresentation analysis was also performed in R, using the Bioconductor package clusterProfiler ^73^. Pearson’s correlation coefficients between HCEC, End1, and HeLa expression of the fibrosis-associated gene set identified by clusterProfiler were subsequently calculated in R, using log_2_-transformed fold changes of all genes differentially expressed in at least one data set. Sequencing data for mock-infected and *Ct* L2-infected HVEs are available in the Gene Expression Omnibus, accession GSE228774; Sequencing data for the equivalent HCEC and End1 infections are also available in GEO as part of our recent publication (GSE180784) ^12^.

### Single-Cell Library Preparation and Sequencing

Infected cocultures were harvested as single-cell suspensions via the 10X Genomics Sample Preparation Demonstrated Protocol (Revision B). Briefly, cocultures were disassociated in 0.25% Trypsin-EDTA, then harvested in complete DMEM. After centrifugation and resuspension in complete DMEM to eliminate residual trypsin, cell suspensions were filtered over 30 um cell strainers (Miltenyi Biotec, 130-098-458) to eliminate clustered cells, washed twice in 1X phosphate-buffered saline (PBS) containing 0.04% bovine serum albumin (BSA), then filtered again using Flowmi 40 uM cell strainers (Bel-Art H13680-0040). After staining aliquots of each suspension with Trypan Blue to validate a minimum of 95% cell viability for each sample, as well as determining suspension density for controlling cell library size. Suspensions were stored on ice for library preparation according to the 10X single-cell protocol (10X Genomics, Chromium Single Cell 3’ v2 Chemistry). cDNA libraries were submitted to the UNMC Genomics Core facility for molarity quantification via qPCR, followed by sequencing using the NextSeq 550 platform and two 150-cycle High-Output flow cells. Sequencing data for this experiment are also available on GEO, accession GSE228647.

### Single-Cell RNA-Sequencing Analysis

Sequencing data was demultiplexed and aligned to the hg38 reference genome via CellRanger (10X Genomics, version 6.0.1), using an expected cell count of 1000 informed by hemacytometer counts performed on an aliquot of the single-cell suspension. After aggregating the mock- and *Ct* L2-infected alignment results for each replicate (n = 2), initial cell type determination was performed in Loupe Cell Browser (10X Genomics, version 3.1.1). Epithelial cell (End1) and fibroblast (KCO2) populations were identified on the basis of keratin 19 and vimentin expression, respectively. Subclustering analysis of the epithelial/fibroblast populations was performed in the *Seurat* R package ^74^. After associating per-cell expression data with their respective treatment (mock- or *Ct* L2-infected) and cell type (epithelial cell or fibroblast) metadata, per-cell expression data were separated into separate End1/KCO2 Seurat objects.

Subsequent quality control and dimensionality reduction was performed analogously to the Seurat guided vignette on single-cell expression analysis. After trimming cells with extreme unique feature counts (200 < n < 8000), outlier features were identified using the vst method (local polynomial regression of log-transformed mean/variance, selecting top 2000 features). The dimensionality of each data set was determined via the Seurat JackStraw method; for both the End1 and KCO2 cell populations, the top 15 principal components encompassed the majority of variance in the data. Dimensionality reduction via uniform manifold approximation and projection (UMAP) was then performed on each data set, using their respective top 15 principal components, followed by cluster identification via Seurat’s shared nearest neighbor algorithm (using a clustering resolution of 0.5). Per-cluster differential expression for each cluster was then determined via the FindMarkers function, using a cutoff of genes expressed by at least 25% of cells in each compared population. Per-cell and per-cluster gene expression was visualized using the Seurat DimPlot, FeaturePlot, and VlnPlot functions, splitting plots across treatment groups or clusters where applicable.

### Immunofluorescence Microscopy

End1s and/or KCO2s seeded on glass cover slips (VWR) and infected at an MOI of 5 as described above were fixed in 4% paraformaldehyde in phosphate-buffered saline (PBS) for 10 minutes at 37° C, washed in PBS, then blocked in 5% bovine serum albumin (BSA) in PBS for 1 hour at room temperature. Fixed and blocked cover slips were subsequently incubated overnight at 4° C with a rabbit anti-collagen I (Abcam ab260043, 1:250 dilution) primary antibody. Cover slips were again washed in PBS, then incubated for 1 hour at room temperature with the following fluorophore-conjugated antibodies/dyes in 1% BSA-PBS: goat anti-rabbit Alexa-488 conjugate (Thermo Fisher Scientific A-11034, 1:1000 dilution), phalloidin Alexa-594 conjugate (Thermo Fisher Scientific A-12381, 1:120 dilution), DAPI (Sigma-Aldrich 10236276001, 1:1000 dilution). Afterward, cover slips were washed in PBS and ultrapure water, then mounted on microscope slides using Shandon Immu-Mount (ThermoFisher Scientific 9990402).

At least 5 fields of each cover slip were imaged using a CSU-W1 Spinning-Disk Confocal Microscope (Nikon). Blinded image quantification was performed by assigning image filenames randomized number codes. For measurement of YAP nuclear translocation, 10 nuclei were selected at random per field using only the DAPI channel, manually masking the nuclear area, and recording the mean YAP fluorescence intensity per nucleus. To account for variation in total YAP between cells, staining efficiency between cover slips, and compression of the nuclear/cytosolic compartments by the chlamydial inclusion, 10 cytosolic regions not occluded by an inclusion body and adjacent to measured nuclei were selected per field, with the mean YAP fluorescence intensity of these regions averaged to produce a per-field measurement of mean cytosolic YAP fluorescence intensity; nuclear translocation of YAP was thereby expressed as a ratio of mean nuclear fluorescence to mean cytosolic fluorescence. For measurement of KCO2 collagen I expression, a selection mask of all cell area per field was defined via fluorescence thresholding of the phalloidin counterstain in ImageJ, per-field measurements of mean collagen I fluorescence intensity were subsequently recorded using this selection mask. Statistical analysis was performed in R, using a Kruskal-Wallis test to first verify a statistically significant (p-value < 0.05) difference between treatment groups. Subsequent pairwise comparisons were performed using a Wilcoxon rank sum test and Bonferroni’s correction for multiple comparisons, with p-values less than 0.05 being considered statistically significant.

### siRNA Transfection

End1s were transfected with either an ON-TARGETplus non-targeting siRNA pool (Horizon Discovery D-001810-10-05) or an ON-TARGETplus YAP1-targeting siRNA SMARTpool (Horizon Discovery L-012200-00-0005) using Lipofectamine 3000 (Thermo Fisher Scientific L3000008), per manufacturer’s instructions, at an empirically determined optimal concentration of 10 nM. At 16 h post-seeding of End1s at 125% of confluence on 6-well transwell inserts as described above, siRNA was combined in Opti-MEM (Thermo Fisher Scientific 31985062) with the Lipofectamine 3000 reagent, incubated for 5 m at room temperature to allow for liposome formation, then added to wells dropwise with mixing. Transfected End1s were then incubated for 24 h prior to infection with *Chlamydia* as described above.

### SDS-PAGE and Western Blotting

To minimize activity of the chlamydial protease CPAF, End1s seeded on 6-well plates, transfected with siRNA, then infected at an MOI of 5 as described above were subsequently lysed in 1% SDS buffer heated to 95° C, as previously described ^75^. After treatment with Pierce Universal Nuclease (Thermo Fisher Scientific, 1:1000 dilution) for 5 minutes at room temperature, lysates were combined with 4X Laemmli Sample Buffer (Bio-Rad 1610747) for loading on a 10% acrylamide/bis-acrylamide gel for SDS-PAGE (1.5 hours, 100V). Gels were then transferred to PVDF membranes (Bio-Rad 1620177) using a semi-dry transfer method (50 minutes, 20V). After blocking in 5% BSA in PBST (PBS containing 0.1% Tween-20) for 1 hour at room temperature, membranes were incubated overnight at 4° C with primary antibodies in 5% BSA-PBST: rabbit anti-YAP (CST 4912, 1:1000 dilution), rabbit anti-GAPDH (CST 2118, 1:1000 dilution). Membranes were subsequently washed in PBST, then incubated with a goat anti-rabbit HRP-conjugated secondary antibody (Dako P0448, 1:2000 dilution in 5% BSA-PBST) for 2 hours at room temperature. After additional washing in PBST, membranes were imaged using Immobilon HRP Substrate (Millipore Sigma WBKLS0500) or an Azure Biosystems c600. Images were analyzed using the ImageJ gel analysis tool to quantify the fluorescence density of total YAP total protein relative to the GAPDH loading control.

### Reverse Transcription Quantitative Real-Time PCR

For measurement of EMT-associated gene expression, End1s were seeded on 6-well plates (Corning 354402) at 100% of confluence and infected at an MOI of 5 as described above. At 24 hpi, RNA was harvested using using TRIzol (Thermo Fisher Scientific 15596026) and the DNA-free DNA removal kit (Thermo Fisher Scientific AM1906), according to manufacturers’ protocols. For measurement of collagen I and α-SMA expression of cocultured fibroblasts, KCO2s seeded on 6-well plates (Corning 354402) and cocultured as described above were harvested for RNA, again using TRIzol and the DNA-free DNA removal kit. cDNA libraries were subsequently prepared using SuperScript IV Reverse Transcriptase (Thermo Fisher Scientific 11766050) according to the manufacturer’s protocol. Quantitative real-time PCR was performed on a QuantStudio 3 (Thermo Fisher Scientific) using TaqMan assay kits (Thermo Fisher Scientific) of the following genes: *CDH1* (Hs01023895_m1), *CDH2* (Hs00983056_m1), *VIM* (Hs00958111_m1), *KRT13* (Hs02558881_s1), *SNAI1* (Hs00195591_m1), *SNAI2* (Hs00161904_m1), *ZEB1* (Hs01566408_m1), *ZEB2* (Hs00207691_m1), *COL1A1*, (Hs00164004_m1), *ACTA2* (Hs00426835_g1), and the housekeeping gene *HPRT* (Hs02800695_m1). Statistical analysis was performed in R, using pairwise Student’s t-tests and Bonferroni’s correction for multiple comparisons; p-values less than 0.05 were considered statistically significant.

## Acknowledgements

The authors acknowledge the members of the Carabeo laboratory for their critical feedback in the development of this research. We also acknowledge the assistance of Maggie Sladek and Megan Otte in performing coculture experiments, Dr. Aileen Helsel and Dr. Nathan Law in single-cell RNA-sequencing experimental design, Bianca Lopez-Biladeau for single-cell RNA library preparation, and the UNMC DNA Sequencing Core in bulk mRNA library preparation and bulk/single-cell RNA-sequencing.

## Author Contributions

Conceptualization, L.C. and R.C.; Methodology, L.C. and R.C.; Validation, L.C.; Formal Analysis, L.C.; Investigation, L.C.; Resources, R.C.; Data Curation, L.C.; Writing – Original Draft, L.C.; Writing – Review & Editing, R.C.; Visualization, L.C.; Supervision, R.C.; Project Administration, R.C.; Funding Acquisition, R.C.

## Data Availability Statement

The bulk RNA-sequencing datasets generated and analyzed for this study can be found in the Gene Expression Omnibus ^95^, GEO Series accession numbers GSE180784 (https://www.ncbi.nlm.nih.gov/geo/query/acc.cgi?acc=GSE180784) (End1 and HCEC bulk RNA-sequencing), GSE228774 (https://www.ncbi.nlm.nih.gov/geo/query/acc.cgi?acc=GSE228774) (HVE bulk RNA-sequencing), and GSE228647 (https://www.ncbi.nlm.nih.gov/geo/query/acc.cgi?acc=GSE228647) (End1/KCO2 single-cell RNA-sequencing).

