## Supplemental Data for "Chlamydial YAP activation in host endocervical epithelial cells mediates pro-fibrotic paracrine stimulation of fibroblasts"

**Supplemental Tables**

**Supplemental Table 1.** Summary of differential gene expression detected in bulk RNA-sequencing of mock- and *Chlamydia trachomatis* serovar L2-infected primary human cervical epithelial cells (HCECs) and human vaginal epithelial cells (HVEs). All fold changes expressed relative to the indicated HVE and/or mock-infected control. See enclosed file: “Supplemental Table 1.xlsx”.

**Supplemental Table 2.** Summary of genes equivalently modulated in an HCEC-intrinsic (mock-infected HCECs vs mock-infected HVEs) and infection-mediated fashion (*Ct* L2-infected HCECs vs mock-infected HCECs). See enclosed file: “Supplemental Table 2.xlsx”.

**Supplemental Table 3.** Summary of transcription factors exhibiting significant enrichment via analysis of the combined HCEC-intrinsic/infection-mediated gene set with ChEA3. See enclosed file: “Supplemental Table 3.xlsx”.

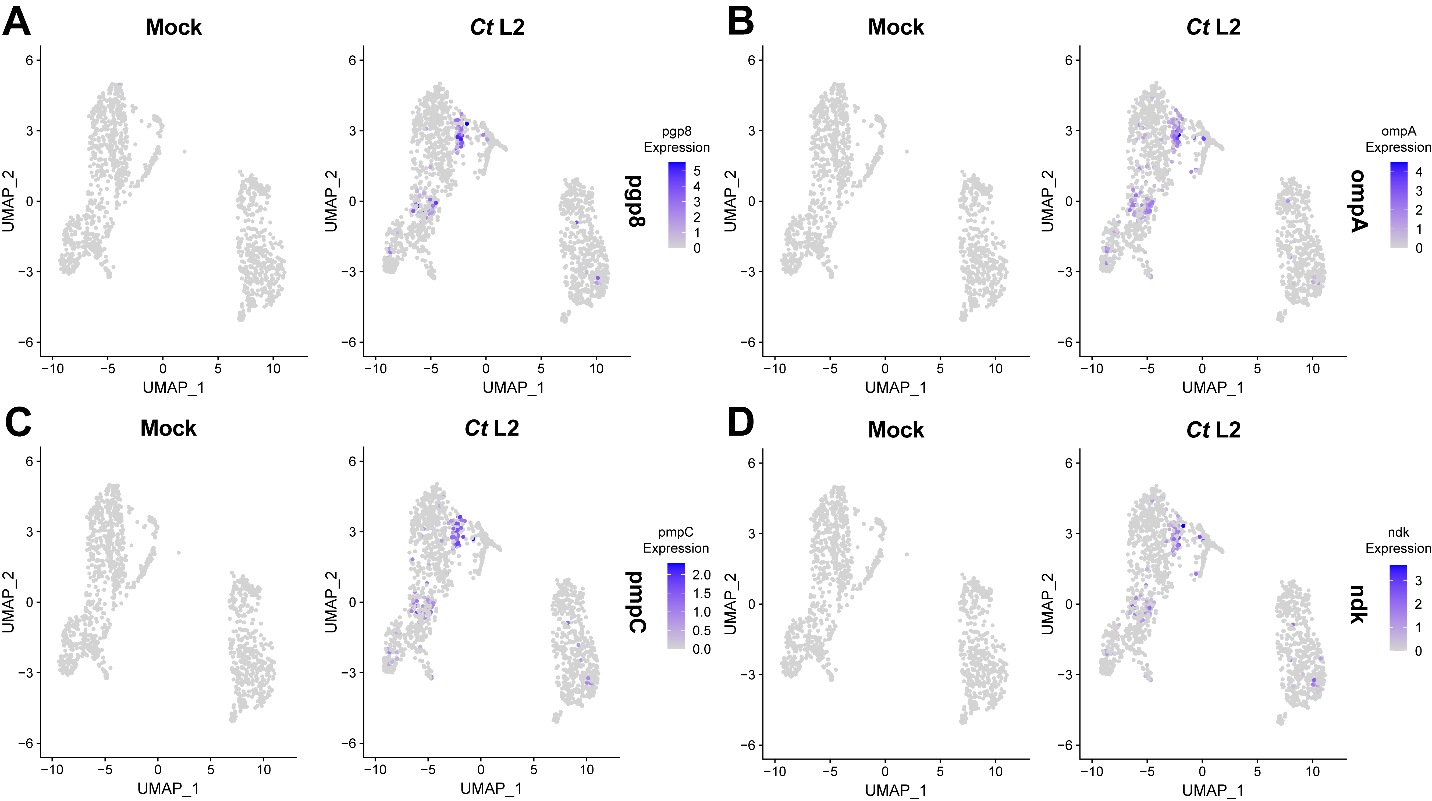
**Supplemental Figures**

**Supplemental Figure 1.** Uniform manifold approximation and projection (UMAP) feature plot of chlamydial pgp8 antisense (**A**), ompA (**B**), pmpC (**C**), and ndk (**D**) expression in same-well coculture of mock- or *Ct* serovar L2-infected End1 endocervical epithelial cells with KCO2 fibroblasts for 23 h (starting at 1 hpi). n = 2 biological replicates, 1000-cell libraries per sample, approx. 5-7 x 10^4^ reads/cell.

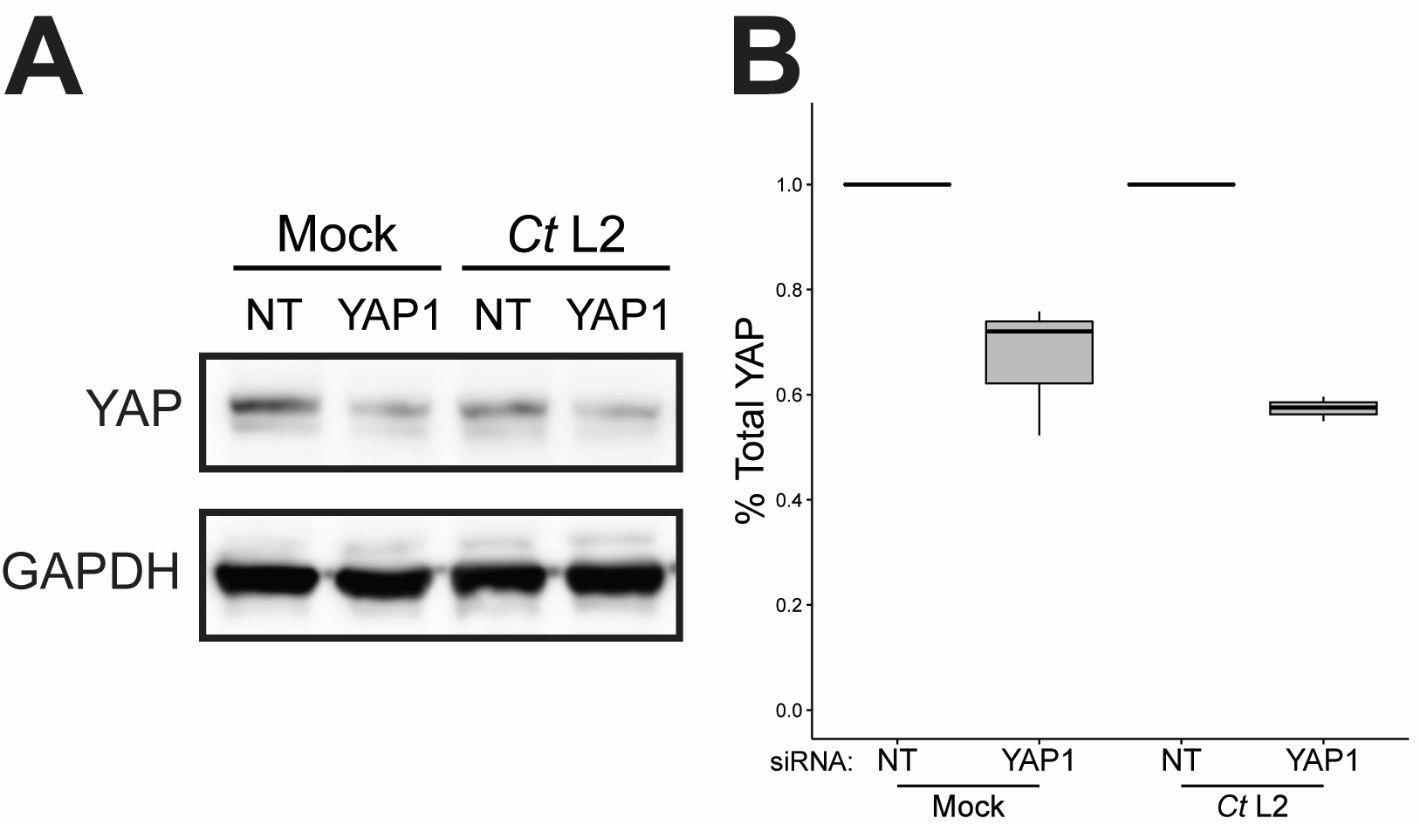
**Supplemental Figure 2.** Efficiency of siRNA-mediated YAP knockdown in *Ct* serovar L2- and mock-infected End1s cocultured on transwells with KCO2s.
(**A**) Representative Western blot of YAP protein and GAPDH loading control in mock- and *Ct* L2-infected End1 cells treated with either non-targeting (NT) or YAP1-targeting siRNA (10 uM for 48 h, starting 24 h prior to infection).
(**B**) Densitometric quantification of total YAP protein after siRNA-mediated knockdown, normalized to GAPDH loading control band. n = 3 biological replicates; whiskers: minimum to maximum; asterisks: p-values ≤ 0.05, using pairwise Student’s t-tests and Bonferroni’s correction for multiple comparisons.
