## Supplementary figures and images for "Chlamydial YAP activation in host endocervical epithelial cells mediates pro-fibrotic paracrine stimulation of fibroblasts"

### Supplemental Figure 1

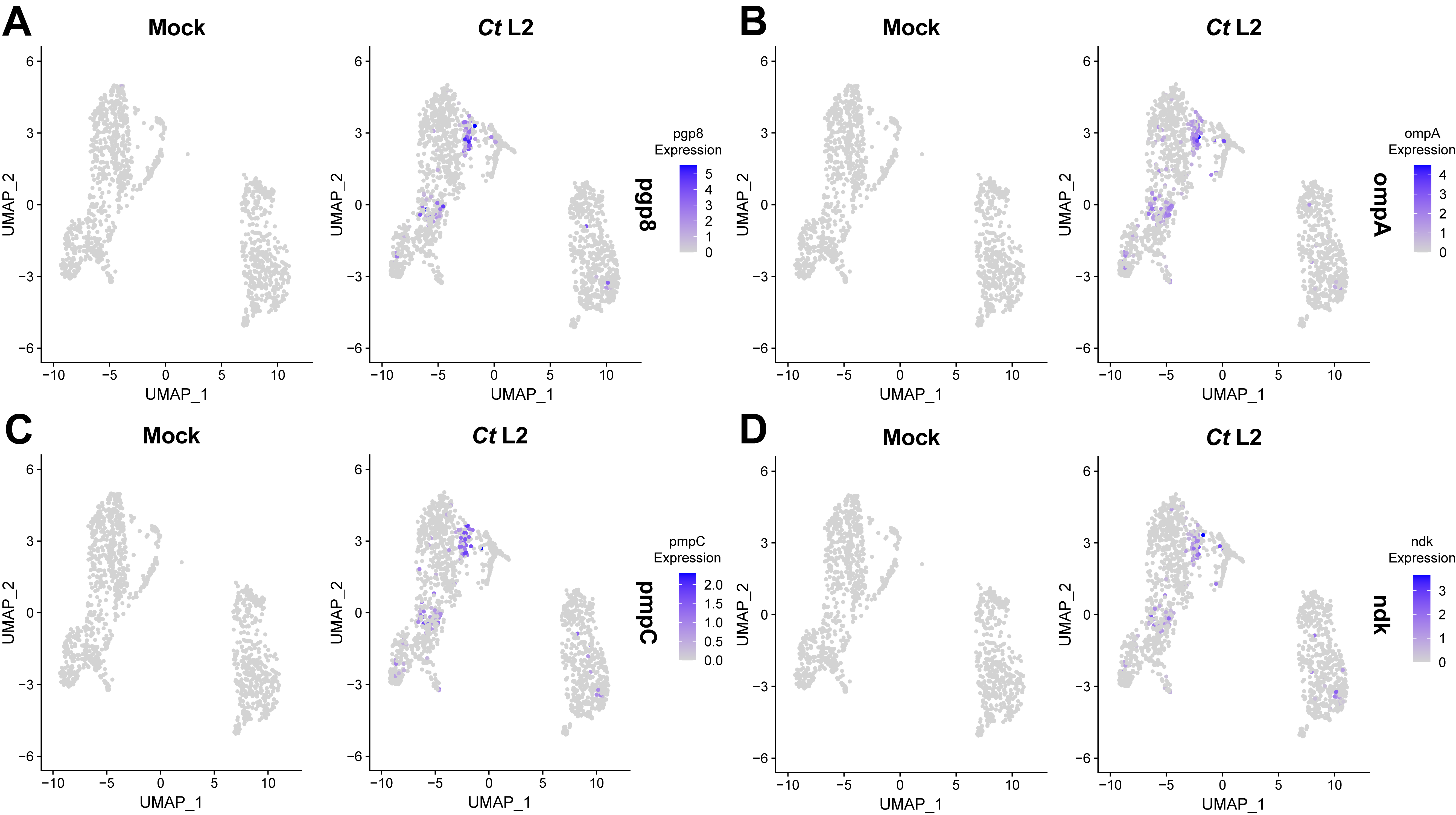

### Supplemental Figure 2

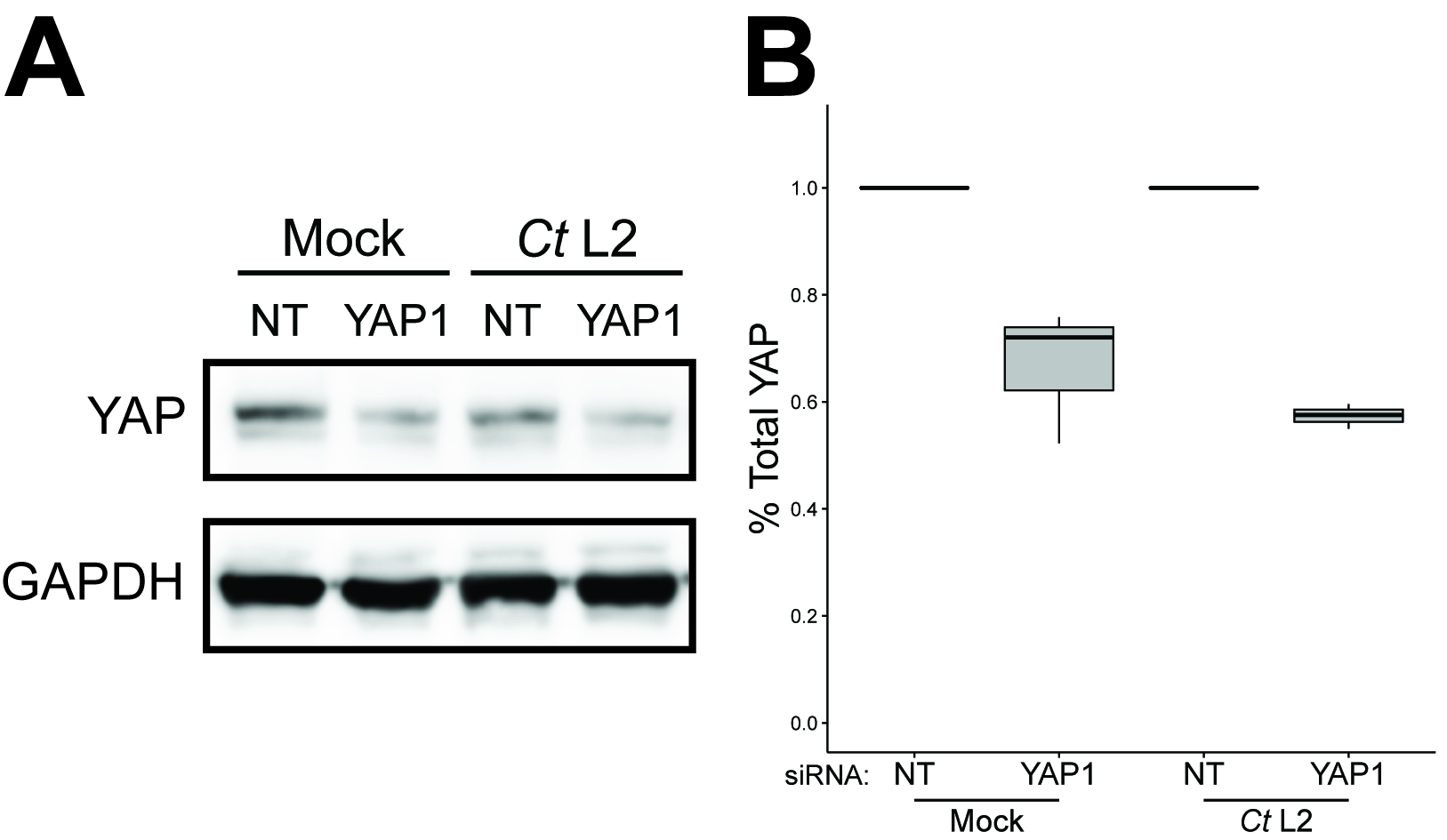
